# Evolutionary chemical learning in dimerization networks

**DOI:** 10.1101/2025.06.16.660005

**Authors:** Alexei V. Tkachenko, Bortolo Matteo Mognetti, Sergei Maslov

## Abstract

We present a novel framework for chemical learning based on Competitive Dimerization Networks (CDNs)— systems in which multiple molecular species, e.g. proteins or DNA/RNA oligomers, reversibly bind to form dimers. We show that these networks can be trained in vitro through directed evolution, enabling the implementation of complex learning tasks such as multiclass classification without digital hardware or explicit parameter tuning. Each molecular species functions analogously to a neuron, with binding affinities acting as tunable synaptic weights. A training protocol involving mutation, selection, and amplification of DNA-based components allows CDNs to robustly discriminate among noisy input patterns. The resulting classifiers exhibit strong output contrast and high mutual information between input and output, especially when guided by a contrast-enhancing loss function. Comparative analysis with in silico gradient descent training reveals closely correlated performance. These results establish CDNs as a promising platform for analog physical computation, bridging synthetic biology and machine learning, and advancing the development of adaptive, energy-efficient molecular computing systems.

**Significance Statement:** This study introduces a new paradigm for learning based on chemical reaction networks rather than digital circuits. Using Competitive Dimerization Networks (CDNs)—biomolecular systems in which species reversibly bind to form dimers—complex classification tasks are learned through in vitro directed evolution. This approach eliminates the need for digital hardware or gradient-based optimization, relying instead on intrinsic molecular dynamics for computation. The resulting chemical classifiers achieve high fidelity and robustness to noise, with performance comparable to that of gradient descent training. These findings establish CDNs as a scalable, energy-efficient platform for molecular computing, suggesting broad potential applications in diagnostics, biosensing, synthetic biology, and nanotechnology, where programmable, adaptive chemical systems could serve as alternatives to conventional electronic processors.

---

Deep Learning is revolutionizing science, technology, and everyday life by enabling artificial intelligence systems to extract patterns and make decisions from vast amounts of data. Traditional approaches rely on digital computation and optimization techniques—such as gradient descent — to train models like Deep Neural Networks (DNNs) (1, 2). However, there is a growing interest in *physical learning*, the notion that physical systems can perform computations through their intrinsic dynamics without relying on conventional digital hardware (3–11). Such approaches harness the response of a properly designed physical system to process information, as demonstrated in optical processors (3, 4), memristive circuits (5), and even mechanical systems (10, 11).

In this context, complex biochemical networks operating inside living cells offer a particularly compelling example of physical learning. In biological organisms, processes such as cellular signaling, gene regulation, and metabolic control exhibit a remarkable ability to respond to environmental cues (12–14). This inherent capacity for complex information processing suggests that the function of evolved biochemical networks can be viewed as a form of *Chemical Learning* (CL), where molecular interactions—governed by thermodynamic and kinetic principles—perform computations analogous to those of artificial neural networks (15, 16).

In this paper, we introduce a CL framework based on the Competitive Dimerization Networks (CDNs)— systems composed of multiple molecular species capable of reversible pairwise binding, where each molecule may form dimers with different partners (17–21). Such systems are ubiquitous in biology, e.g. receptor–ligand binding networks, pairwise interactions between transcriptional factors used in combinatorial gene regulation, etc. The computational potential of proteinbased synthetic dimerization networks has been explored in Refs. (19, 20). More recently, engineered CDNs based on non-complementary DNA oligomers have been proposed as a novel platform for DNA computing (22). Recently it was shown that combining DNA hybridization with covalent binding can be used to construct DNA-based CDNs that convert multi-channel inputs into controlled modulation of specific dimer concentrations (21). We anticipate that the extended use of synthetic CDNs based on proteins, DNAs, or RNAs will become increasingly prevalent.

Some of the physical learning proposals require *in silico* training—that is, computer simulations to determine optimal parameters via methods such as backpropagation and gradient descent; others allow for *in-situ* training (7–9). Similarly there have been proposals of employing in silico machine-learning tools to design chemical and biological networks with desired functionality (23, 24). A key feature of our proposed CL framework is the possibility of *in vitro* training via directed evolution. In this approach, the desired function is acquired by the CDN without prior knowledge of its parameters or the need for precise engineering of the optimal system. Instead, we outline a protocol through which the optimal realization is obtained via artificial selection over a series of trials.

## Model and Protocols

### Competitive Dimerization Networks

In our model, a **competitive dimerization network (CDN)** consists of *N* distinct molecular species (e.g., DNAs or proteins) that can form pairwise dimers (see Fig. 1a). The formation of complexes larger than dimers is assumed to be negligible. We denote the overall concentrations of these molecules by **c** = (*c*_1_*, c*_2_*, …, c_N_*) and their fugacities––the fractions of total concentration that remain unbound––by **f** = (*f*_1_*, f*_2_*, …, f_N_*).

**Fig. 1.**
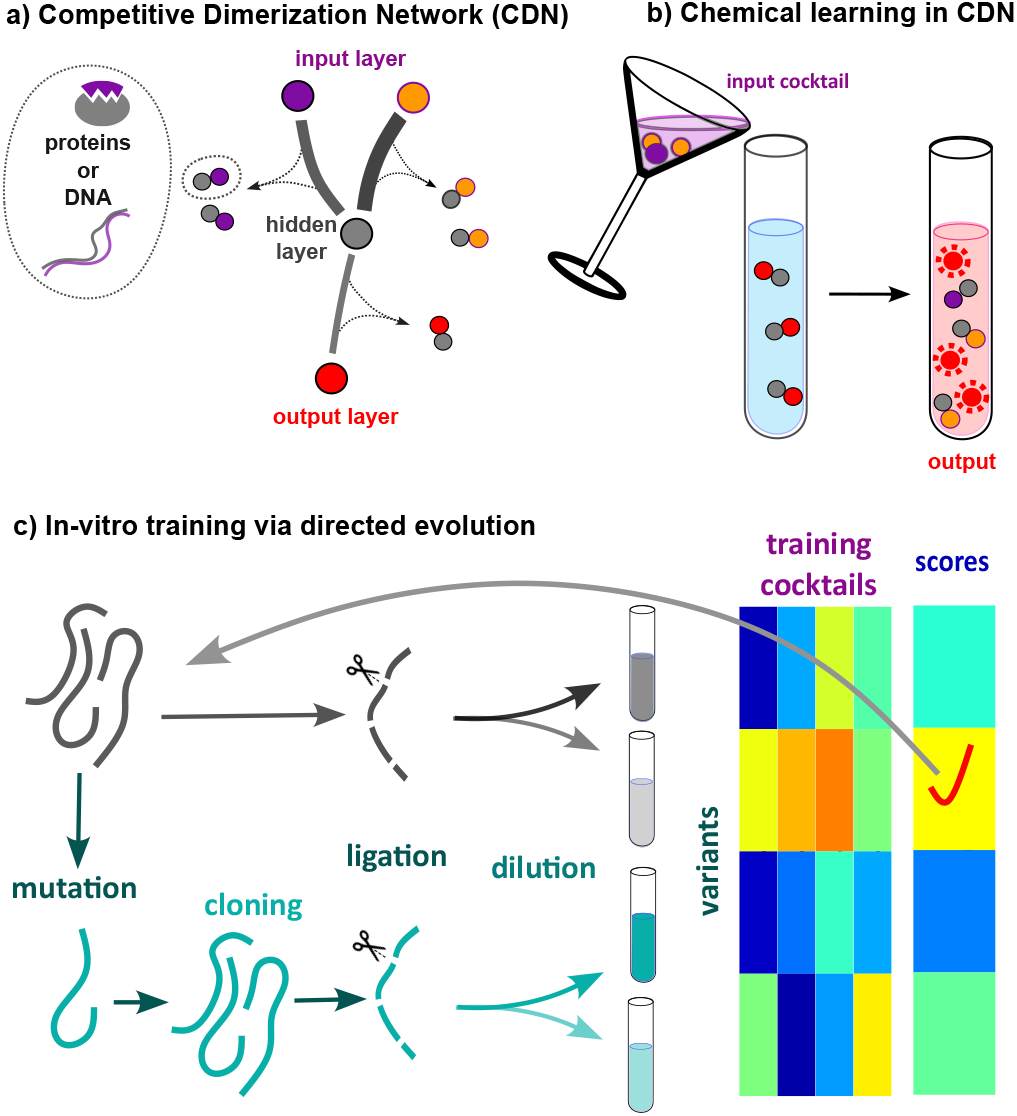
Overview of chemical learning in competitive dimerization networks. (a) Schematic of a network composed of molecular species that reversibly form pairwise dimers, with designated input, hidden, and output layers. (b) Illustration of chemical computation: input cocktails modulate mass action binding equilibria to generate output fugacities. (c) In vitro training protocol using directed evolution, involving mutation, ligation, dilution, and selection of DNA sequences and their concentrations to optimize classification performance.

In chemical mass-action equilibrium free concentrations (activities) of the molecules, *x_i_*= *f_i_c_i_*must satisfy the following set of equations:

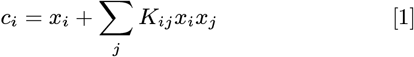

Here, the symmetric association constant *K_ij_*= *K_ji_*characterizes the strength of dimer formation between molecules of types *i* and *j*. The above equilibrium conditions yield the following equations for fugacities *f_i_*:

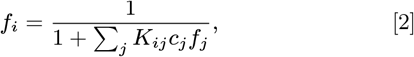

Despite its relative simplicity, we demonstrate below that the CDN is capable of performing learning tasks analogous to those carried out by artificial neural networks. Moreover, we show that the chemical network can be trained *in vitro* using directed evolution. In our framework, each molecular species functions as a “neuron,” and the dimensionless coefficients *K_ij_c_j_*play the role of “synaptic weights” connecting these neurons. Because these weights depend on both the concentrations *c_i_*and the association constants *K_ij_*, they can, in principle, be experimentally tuned by adjusting the composition of the mixture or by modifying (or mutating) the molecules.

In analogy with DNNs, we designate: (i) a subset of *n_in_*molecular concentrations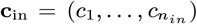 as the input layer; (ii) a non-overlapping subset of *n_out_*fugacities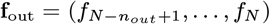asthe output layer; and (iii) the remaining *n_h_* = *N* − n_*in*_− *n_out_* molecules constitute a hidden layer (see Fig. 1a).

Training the CDN involves adjusting both the concentrations (outside of the input layer) and the association constants to achieve a desired functionality. Once trained, the system’s output for any given input can be determined by measuring the output fugacities––for example, via FRET labeling. Because the input is represented by a set of concentrations of species mixed together, we refer to it as an “input cocktail” (see Fig. 1b). Numerically, the simplest way to compute the model’s output is to iteratively solve the mass-action equilibrium equations 2 for **f** .

Although our system bears conceptual similarities to DNNs, several important distinctions must be noted. First, the network structure of the CDN is inherently bidirectional––in contrast to the typical feedforward architecture of DNNs––since its connections, determined by the association constants, are naturally reciprocal. Second, the weights in the CDN (i.e., the coefficients *w_ij_*) are strictly nonnegative. Third, the nonlinear activation function in the equilibrium equation,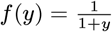, emerges directly from the law of mass action and may not be optimal for all learning tasks.

Notably, because our model corresponds to the minimization of a well-defined free energy, it shares similarities with Hopfield neural networks (25, 26)––one of the simplest realizations of associative memory.

### In-vitro Training via Directed Evolution

While the generic description of CL in CDNs supports multiple molecular implementations, one of the simplest realizations is based on noncomplementary DNA oligomers (22). In this approach, the binding affinities *K_ij_*are determined by the specific oligomer sequences and can be readily modulated via mutations.

We propose a concrete experimental implementation for the *in vitro* training of such CDNs, as illustrated in Fig. 1c. In our design, the sequences corresponding to molecules in the hidden and output layers are concatenated into one or several *master sequences* that serve as the system’s genetic code. At each round of directed evolution, these master sequences are replicated with randomly induced mutations. Each master sequence is PCR-amplified and purified by removing complementary components generated by the PCR (e.g., using magnetic beads). Subsequently, the master sequences are cleaved into their constituent DNA oligomers (for instance, using restriction enzymes), diluted to desired concentrations *c_i_*, and mixed with a cocktail of input oligomers at specified concentrations, **c**_in_. The output fugacities, **f**_out_, are then detected using techniques such as FRET.

Training the system involves exposing several CDN variants to batches of input cocktails. These variants are ranked *in silico* according to a loss function that quantifies the similarity between the measured and desired output fugacities. The best-performing variant is selected for the next round of *in vitro* evolution (see Fig. 1c). Differences among variants arise both from mutations that alter the association constants *K_ij_*and from variations in the composition of the hidden and output layers, as determined by the concentrations of the master sequences.

In the present study, we model the effect of mutations as a biased multiplicative random walk of the association constants *K_ij_*:

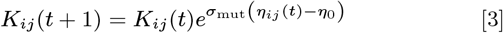

Here, *t* denotes the round of in vitro evolution, and *η_ij_*(*t*) is a Gaussian white noise with *η_ij_*(*t*) = 0 and ⟨*η_ij_*(*t*)*η_ij_*(*t^1^*)⟩ = *δ_tt′_* . The variance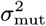characterizes the mutation rate and its impact on binding free energy, while the drift parameter *η*_0_*>* 0 reflects the mutational bias toward weakening binding interactions. Note that (i) ln *K_ij_*is proportional to the binding free energy between molecules *i* and *j*, which justifies our choice of a multiplicative random walk; and (ii) mutations are statistically more likely to weaken binding rather than strengthen it, rendering the random walk negatively biased. In a more detailed study, one may explicitly introduce random mutations into the sequences of individual DNA oligomers and calculate *K_ij_*using the standard model described in (27).

### Chemical Classifier

While CDNs can, in principle, perform a wide variety of learning tasks, here we demonstrate their utility as a multiclass classifier. In our study, a certain number of different of classes of input cocktails are generated randomly, as shown in Fig. 2a. Each cocktail is characterized by a set of input concentrations, **c**_in_, drawn from a log-normal distribution. To emulate the noise present in real experiments, these input concentrations are further scrambled by multiplicative noise. For the outputs, one-hot encoding is employed so that the number of outputs, *n_out_*, equals the number of cocktail classes to be distinguished. As illustrated in Fig. 2b, one-hot encoding designates a specific output as active (high fugacity or “on”) for each class, with all other outputs remaining inactive (low fugacity or “off”).

**Fig. 2.**
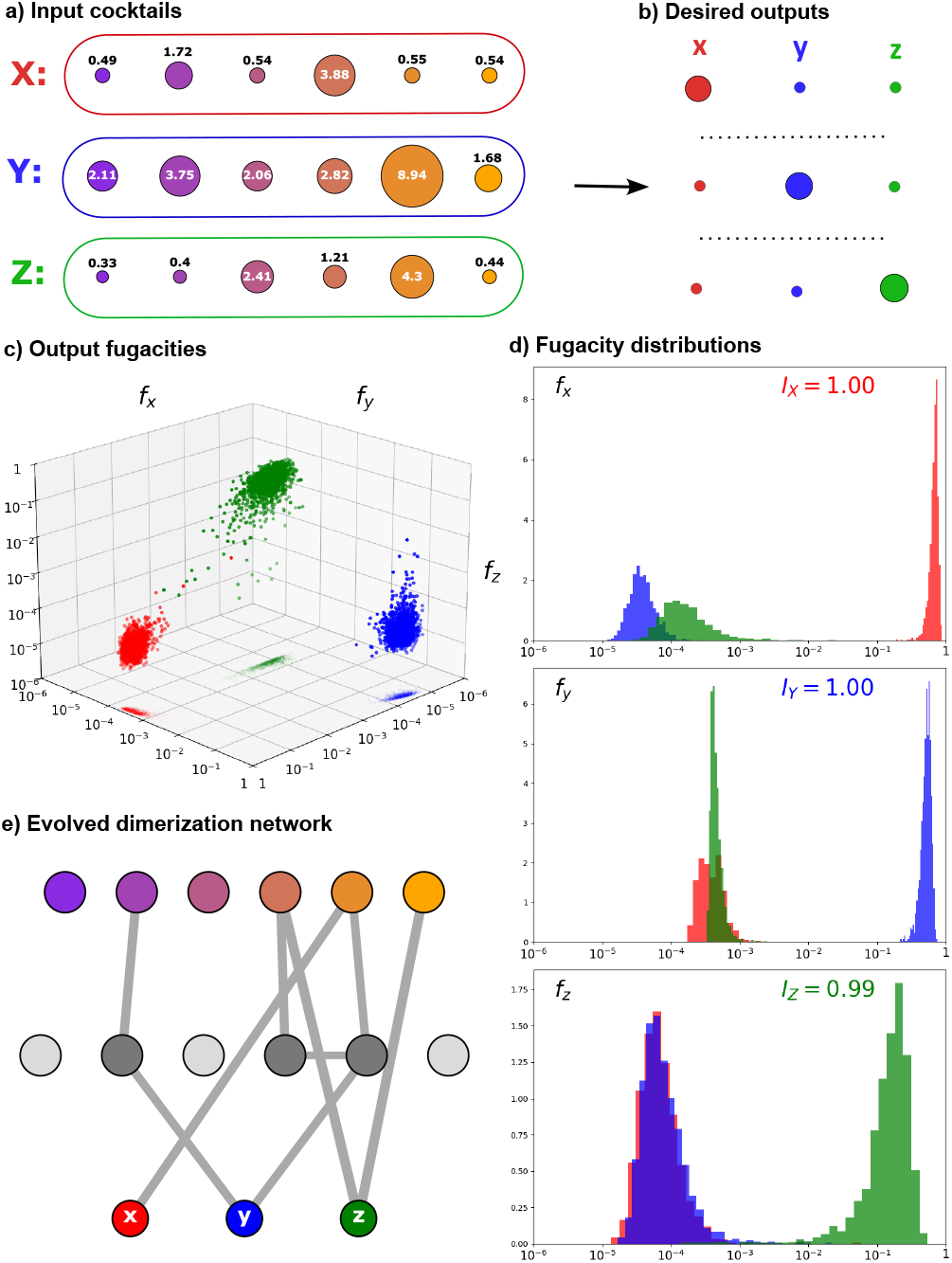
Performance of the chemical classifier trained via directed evolution. (a) Randomly generated three input cocktails. (b) One-hot encoded target outputs for three cocktail classes (X, Y, Z). (c) 3D scatter plot showing class separation based on output fugacities at 20% noise. Each point corresponds to a single noisy realization of inputs in the test dataset. The colors correspond to the true class of the cocktail. (d) Histograms of output fugacities plotted in panel c, showing strong contrast between “on” and “off” states. (e) Network diagram of the evolved CDN after 400 generations of in vitro evolution. Grayscale shading of hidden layer nodes indicates their overall connectivity. In this specific example, all displayed interactions have identical association constants, fixed at the upper saturation limit. Interactive version of this figure, allowing scanning over multiple parameters and cocktail sets, is available at: https://atkachen00.github.io/ChemClassifierWeb

At each step of in vitro evolution, we generate a balanced batch in which each class of patterns is represented by an equal number of examples and each example subject to a different realization of noise. Unlike traditional deep neural network training—which typically involves gradient descent and back-propagation—our numerical experiments utilize an evolutionary learning process. In this process, variants are generated by (i) randomly and independently mutating all association constants according to Eq. 3, and (ii) modulating the overall concentrations of molecules in the hidden and output layers as illustrated in Fig. 1c. Experimentally, this setup can be realized by first amplifying batches with mutations introduced via random mutagenesis using PCR and then diluting them by preset factors. The winning variant, characterized by a specific combination of sequences and concentrations, is selected based on a custom loss function.

Since the goal of our training is to maximize the contrast between the “on” and “off” states of the output components, the loss function is designed to penalize poor contrast. For each output component, we compute the ratio of the *arithmetic mean* of its “off” concentrations to the *geometric mean* of its “on” concentrations. The overall loss is defined as the logarithm of the worst (i.e., highest) contrast ratio across all output components:

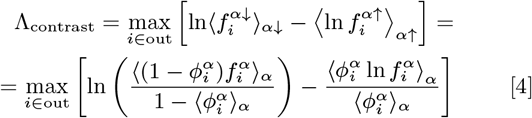

In this formulation, the index *α* labels the samples in the training batch, and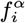represents the output fugacity for component *i* in sample *α*. The variable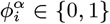denotes the desired target state of component *i* for sample *α*. The subsets of the batch where component *i* is expected to be “on” or “off” are denoted by *α*↑ and *α*↓, respectively (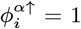and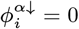). The notation *⟨·*⟩↑*_α_* indicates an average over the full batch, while ⟨*·*⟩*_α_*↑ and ⟨*·*⟩↓*_α_* denote averages taken over the “on” and “off” subsets for each output component *i*, respectively. The maximum function selects the worst performing (the least negative) output component *i*. For the loss function to be applicable, it is necessary that 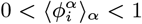for each output component *i*, ensuring that both “on” and “off” samples are present in the batch.

## Results

### Performance and Efficiency of the Chemical Classifier

We numerically investigated a chemical classifier comprising *n_out_*= 3 classes of cocktails (labeled *X, Y*, and *Z*) with *n_in_*= 6 input channels. The input class concentrations *c_i_*were randomly generated from a log-normal distribution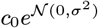with *σ*^2^ = 2. The actual concentrations presented to the chemical classifier were then scrambled by the multiplicative noise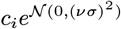, emulating inevitable noise present in real experimental conditions. The parameter *ν* thus quantifies the noise-to-signal (N/S) ratio in our inputs. Figure 2 illustrates one realization of inputs generated with a N/S ratio set to 20%. At this noise level, the evolutionarily trained CDN successfully discriminates among the three classes. This is evident in the 3D scatter plot (Fig. 2c), where data points form three distinct clusters—red for *X*, blue for *Y*, and green for *Z*.

Furthermore, the fugacities of individual output components (*x, y*, and *z*) clearly resolve the classes, as demonstrated by the histograms in Fig. 2d. The contrast between the output fugacities of the “on” and “off” states spans three orders of magnitude. Note that all axes in Figs. 2c–d are displayed on a logarithmic scale.

Figure 2e presents the structure of the evolutionarily trained chemical network after 400 generations. In this network diagram, the thickness of each edge reflects the logarithm of the corresponding association constant *K_ij_*. For clarity, edges with *K_ij_<* 1 have been omitted since their contributions to the classifier function are negligible. The strongest connection corresponds to *K_ij_*= 10^4^, which is the upper cutoff set in our evolution procedure.

The efficiency of the chemical classifier is evaluated using two complementary metrics. The first measure, the on/off contrast, is defined as the logarithmic difference (expressed in decades) between the output fugacities in their “on” and “off” states. The second metric is the Mutual Information (MI) between the inputs and outputs, an information-theoretic quantity that generalizes the correlation coefficient. In this context, MI captures the degree of overlap between the “on” and “off” histograms of the outputs.

The mutual information between the binary variable 𝒳(assigned a value of 1 for cocktails of class *X* and 0 otherwise) and the corresponding output fugacity *f_x_*is given by:

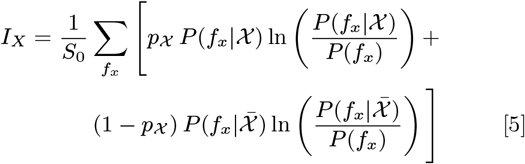

Here, *p*_𝒳_= 1*/n_out_* denotes the fraction of samples corresponding to class *X*, and

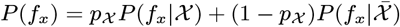

is the total probability of observing a particular fugacity *f_x_*. This expression assumes a balanced testing set, with equal representation of each of the *n_out_*classes. The normalization factor *S*_0_− = (*p*𝒳 ln *p*𝒳 + (1 − p𝒳 ) ln(1 − p𝒳 )) is selected so that *I_X_* = 1 corresponds to the maximal MI—achieved when the “on” and “off” histograms of *f_x_*are completely nonoverlapping—while *I_X_*= 0 indicates statistical independence between 𝒳 and *f_x_*. Analogous definitions hold for the outputs *y* and *z*, with the high fidelity of our chemical classifier evidenced by values *I_X_, I_Y_*, and *I_Z_*all approaching 1 (Fig. 2d).

### Effects of noise and training parameters

Figure 3 illustrates how both the noise level and the choice of loss function affect the efficiency of our classifier. As expected, an increase in the noise-to-signal (N/S) ratio, *ν*, results in a deterioration of performance (Figs. 3a–d). Specifically, comparing panels (a) and (b) for an N/S ratio of 40% with panels (c) and (d) for 100% clearly shows that MI drops markedly as noise increases, evident from the increased overlap between the “on” (red) and “off” (blue and green) histograms.

**Fig. 3.**
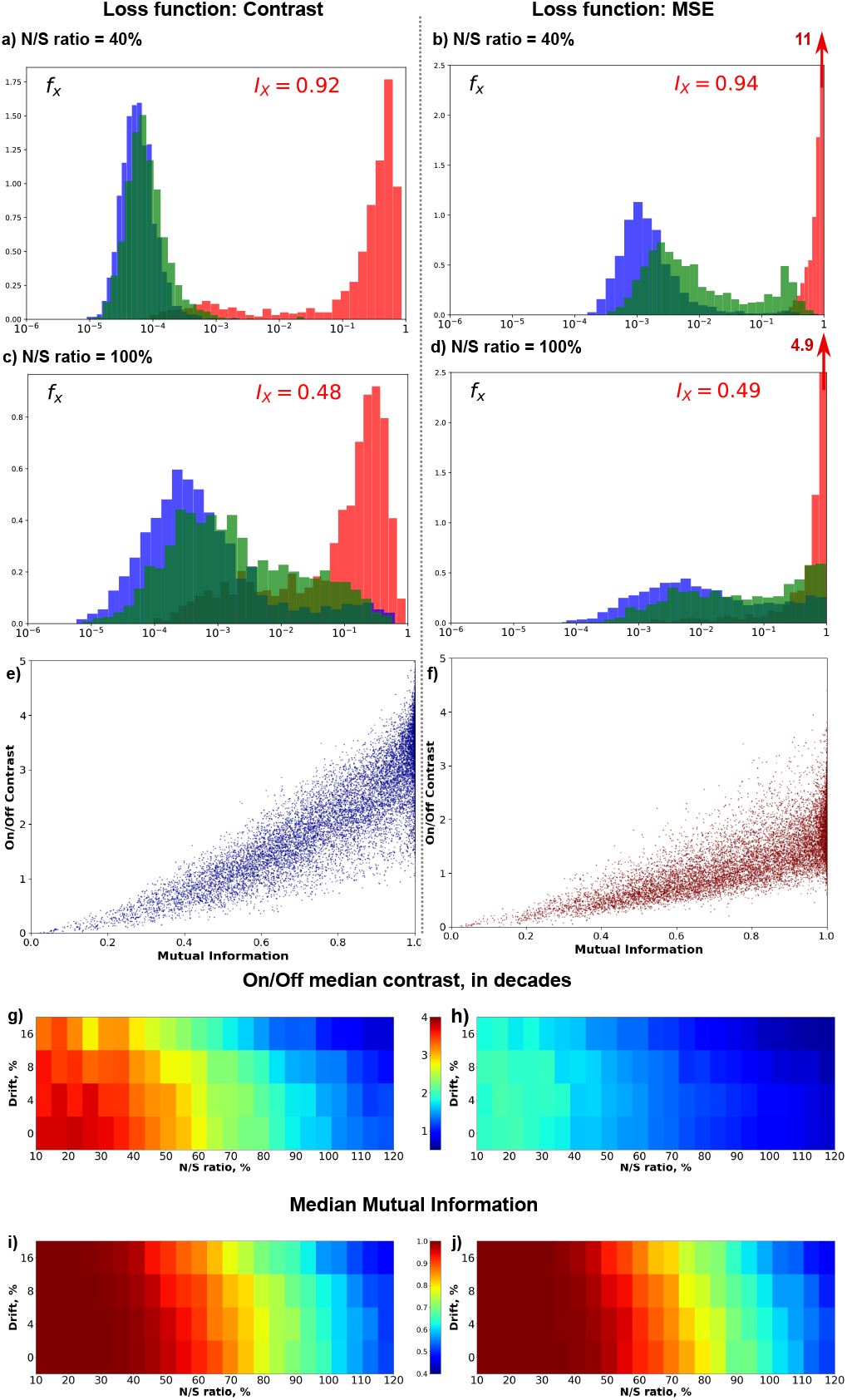
Effect of noise and choice of loss function on classifier performance. Throughout the figure, the left column corresponds to the contrast loss, while the right column corresponds to the MSE loss. (a–d) Output histograms and mutual information at 40% and 100% noise-to-signal ratios, showing the effect of the loss function on output separation and information content. (e, f) Scatterplots illustrating the correlation between on/off contrast and mutual information for both loss functions. Points correspond to different realizations of input classes, noise levels and drift parameters. (g, h) Heatmaps of on/off contrast as a function of drift and noise-to-signal ratio. (i, j) Corresponding heatmaps of mutual information, demonstrating the robustness of information transmission under varying training conditions.

The selection of the loss function plays a critical role, particularly in terms of On/Off contrast. Our bespoke loss function, Eq. 4, used for generating Figs. 3(a) and (c)—is specifically engineered to maximize contrast, yielding significantly better on/off separation than the conventional mean-squared error (MSE) loss function, which was employed in Figs. 3(b) and (d). While the MSE loss produces a sharper peak in the “on” state, it results in a weaker On/Off contrast. These findings indicate that the optimal choice of loss function should be guided by the specific requirements of the intended application. The role played by the choice of the loss function, and the connection between the two metrics used, is apparent from the comparison of two scatter plots, 3(e) and (f). These plots indicate a strong correlation between the Mutual Information, and On/Off contrast parameter, but the latter provides a valuable additional insight once MI saturates at its maximum value 1. As we have already observed in the specific example above, the use of MSE loss function results in significantly lower On/Off contrasts.

Heatmaps in Figs. 3g–j further quantify these trends as functions of both the noise-to-signal (N/S) ratio (x-axis) and the negative drift parameter *η*_0_from the evolutionary training defined in Eq. 3 (y-axis). Increases in either the noise-to-signal ratio or the drift parameter degrade classifier performance. Notably, while the loss functions Λ_contr_(Fig. 3(g) and MSE (Fig. 3(h) exhibit dramatically different on/off contrasts, the mutual information remains virtually unchanged, as shown in Figs. 3(i)–(j).

Increasing the drift parameter leads to a modest decrease in the chemical classifier’s noise tolerance while simultaneously rendering the trained dimerization network sparser. In other words, the network achieves its function through a smaller number of strong links, as demonstrated in Figs. 4ab. To construct these graphs, we disregarded all weak interactions satisfying *K_ij_c*_0_*<* 1. The thickness of the remaining edges is proportional to ln *K_ij_c*_0_, which, in turn, is proportional to the corresponding binding free energies. Specifically, the edge weights *w_ij_*are defined as

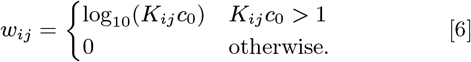

**Fig. 4.**
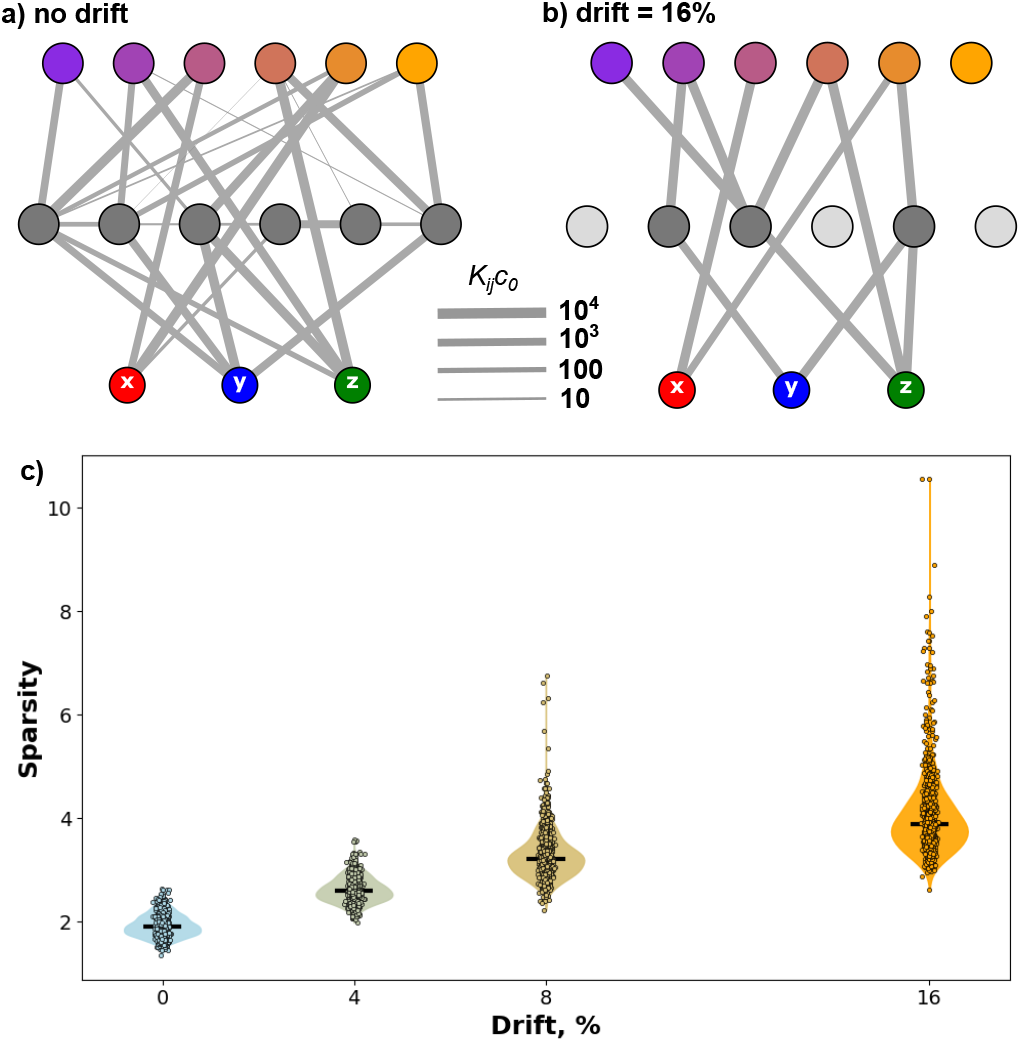
Drift-induced sparsity in trained chemical networks. (a, b) Evolved CDN network topologies at zero drift (left) and 16% drift (right), with edge thickness proportional to log-transformed binding strengths (*K*_0_*c*_0_). Weak interactions (*K*_0_*c*_0_*<* 1) are omitted. Grayscale of hidden nodes reflects their connectivity, defined as the sum of their edge weights. (c) Network sparsity measured by the dispersion coefficient (standard deviation divided by mean) of edge weights as a function of the evolutionary drift.

We quantify the sparsity of a weighted network in terms of the dispersion coefficient (the ratio of the standard deviation to the mean) of its edge weights *w_ij_*, and we show the relationship between sparsity and the evolutionary drift parameter in Fig. 4c. The observed increase in sparsity with drift is expected, as the drift tends to weaken all bindings and eliminate redundant interactions. In this regard, the drift plays a role analogous to regularization in standard machine learning. Consequently, it is not surprising that the simpler, sparser networks exhibit lower noise tolerance compared to their more connected counterparts (see Figs. 3g–h).

### Comparison between Evolutionary Learning and Gradient Descent

All of the results discussed above were obtained using evolutionary training of the chemical classifier, a method that can be implemented in vitro. In contrast, a more traditional approach to training neural networks relies on backpropagation and Gradient Descent (GD). However, because our system is bidirectional—unlike standard feedforward neural networks—backpropagation cannot be applied. Nonetheless, GD remains feasible and is implemented as described in the SI Appendix. As noted earlier, the regularization coefficient in the GD algorithm serves a role analogous to the drift parameter in Evolutionary Learning (EL). It is therefore instructive to compare CDNs trained using EL and GD (see Fig. 5). Each point in the scatterplot represents a specific combination of input cocktail and noise level. Because there is no one-to-one mapping between the drift parameter and the regularization coefficient, each point reflects an average over multiple values of drift (for EL) and regularization (for GD). As shown in Fig. 5a-b, the performance of the two training methods is highly correlated and generally comparable. However, GD exhibits a slight advantage over EL, as indicated by the clustering of points below the diagonal dashed line in both panels.

**Fig. 5.**
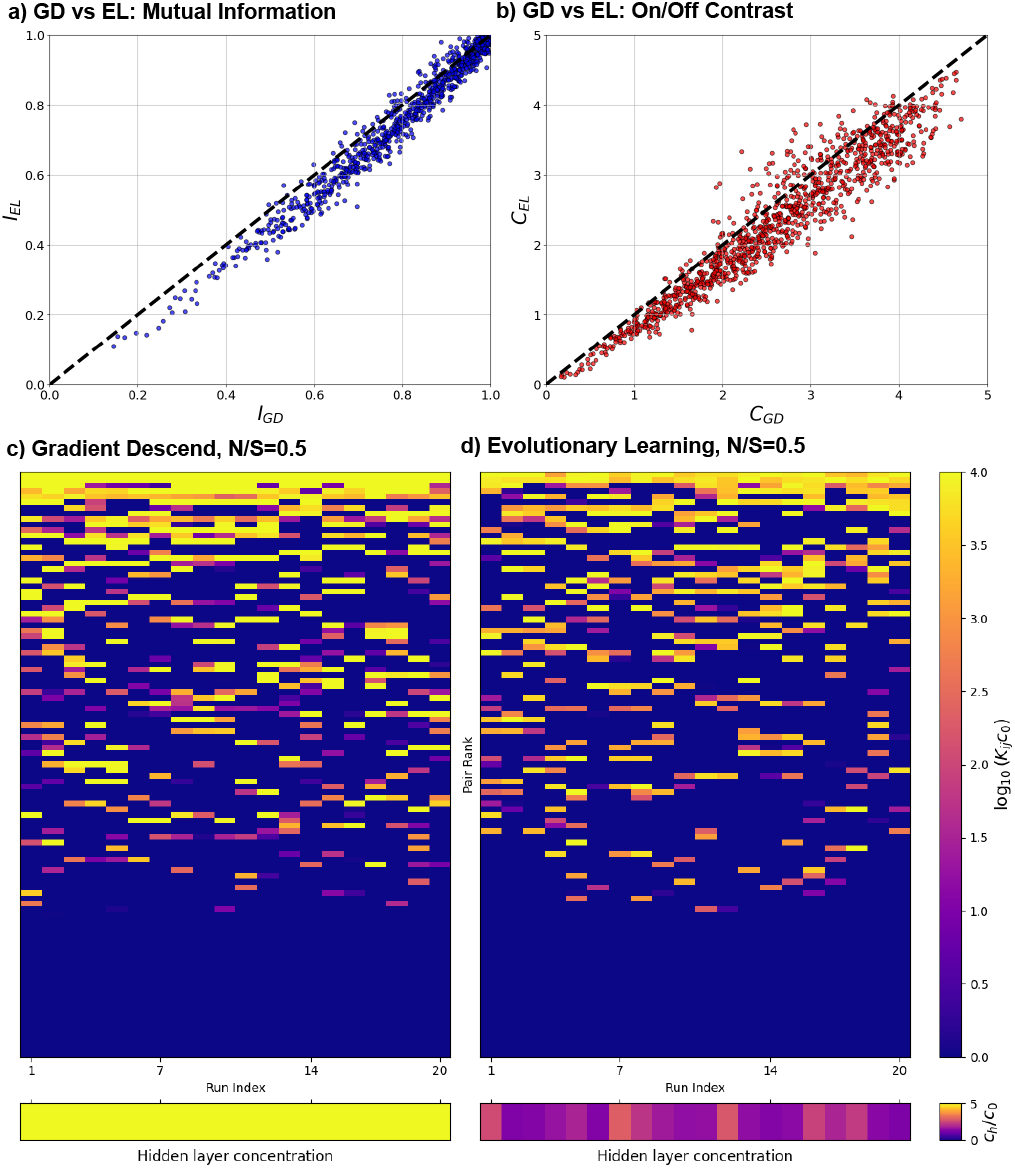
Comparison of network structures obtained by evolutionary learning (EL) and gradient descent (GD). **(a,b)** Scatter plots comparing the chemical classifier performance for EL and GD across multiple input cocktails and noise levels. Panel (a) shows mutual information, and panel (b) shows on/off contrast. Each point represents performance averaged over multiple drift (EL) and regularization (GD) parameters. The dashed diagonal indicates equal performance; points below the diagonal correspond to cases where GD outperforms EL. **(c,d)** Heatmaps summarizing network variability across 20 independent training runs for GD (panel c) and EL (panel d), performed on the same cocktail set at *N/S* = 0.5. The top sub-panels show edge weights *w_ij_* = log_10_(*c*_0_*K_ij_* )*, c*_0_*K_ij_>* 1, ordered by their mean strength across all runs; the bottom sub-panels show hidden layer concentrations *c_h_/c*_0_. Columns correspond to individual training runs and are hierarchically clustered by network similarity.

To further probe the distinction between GD and EL, we compared the network structures and their reproducibility under each training scheme. We performed 20 independent runs of both EL and GD on the same cocktail set at *N/S* = 0.5. Figure 5c–d presents these results as heatmaps of edge weights *w_ij_* and hidden-layer concentrations *c_h_/c*_0_. Rows correspond to 90 pairs of interacting molecules, ranked by their interaction strength ⟨*w_ij_*⟩ averaged over all runs, while columns represent individual runs. Strikingly, none of the runs produced an identical network topology, underscoring a highly degenerate and rugged loss-landscape. Despite this diversity, classifier performance remains consistent across runs. In GD, five of the strongest network links persist across most solutions, and the hidden-layer concentration invariably reaches its maximum, *c_h_* = 5*c*_0_. In contrast, EL retains only two invariant links and exhibits considerable run-to-run variability in *c_h_*, which never attains the upper bound. From this perspective, EL behaves as though sampling at a higher effective temperature compared to GD. Remarkably, despite these significant structural differences, the final classifier performance under both methods remains nearly identical (Fig. 5a–b).

## Discussion and Conclusions

In this study, we developed and explored the concept of chemical learning in competitive dimerization networks (CDNs) and demonstrated their in vitro training through directed evolution. Our results establish that CDNs, which harness intrinsic biomolecular interactions, are capable of performing complex classification tasks with high fidelity and robust noise tolerance.

We found that our chemical classifier can reliably discriminate among various input cocktails by employing a one-hot encoding strategy. In this scheme, each cocktail triggers the activation of a specific output component by transitioning it to a free, unbound state, while all other outputs remain predominantly in a tightly bound state. This clear separation results in a substantial on/off contrast, with measurable differences that span several orders of magnitude.

A key aspect of our work is the use of in vitro training via directed evolution. In contrast to traditional deep learning approaches that depend on gradient descent and backpropagation, our protocol enables the network to iteratively evolve toward the desired functionality through selective pressure. This evolutionary process not only optimizes the association constants and concentrations within the network but also naturally regularizes the system by progressively eliminating redundant interactions, leading to sparser and more interpretable networks.

Furthermore, the possibility of a hybrid training strategy is particularly promising. An initial in silico phase can be used to design an approximate optimal dimerization network by determining target association constants through computational simulations. These computationally derived parameters can then be implemented in the physical system by engineering appropriate intramolecular interactions—such as selecting the right sequences of DNA, RNA, or proteins—and subsequently refined with in vitro directed evolution. This approach effectively marries the speed and flexibility of digital simulation with the adaptability and robustness intrinsic to experimental evolution.

The broader implications of our work extend well beyond the proof-of-concept demonstrated here. High-fidelity CDNs may find applications across a range of fields, from biomedical diagnostics—where they could help distinguish between healthy and diseased cellular states—to nanotechnology, in which adaptive chemical networks might mediate environment-responsive structural transformations. These potential applications highlight the versatility of chemical networks as a platform for unconventional information processing.

Overall, our study underscores the promise of chemical learning as an innovative paradigm for computation. By leveraging the laws of thermodynamics and kinetics inherent to biomolecular systems, CDNs offer a novel approach to information processing that sidesteps the need for conventional digital hardware. Inspired by evolution and intermolecular interactions in biological systems, this work lays the foundation for future advances in adaptive and energy-efficient molecular computing.

## ACKNOWLEDGMENTS

This research used the Theory and Computation facility of the Center for Functional Nanomaterials (CFN), which is a U.S. Department of Energy Office of Science User Facility, at Brookhaven National Laboratory under Contract No. DE-SC0012704. Computational resources have also been provided by the Consortium des Équipements de Calcul Intensif (CECI), funded by the Fonds de la Recherche Scientifique de Belgique (F.R.S.-FNRS) under Grant No. 2.5020.11 and by the Walloon Region. Codes and data generated as a part of this work are available via GitHub repository https://github.com/atkachen00/ChemClassifierWeb

## SI Appendix

### A. Loss functions

In our modeling of evolutionary training we employed two different loss functions, one of which is the “contrast” loss introduced by Eq.(4) in the main text, the other is the standard Mean Square Error (MSE) between the desired and observed output fugacities:

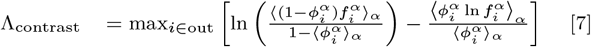

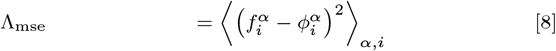

### B. Performance metrics

The performance of the chemical classified was characterized by using the Mutual Information, Eq. 5, and on/off Contrast:

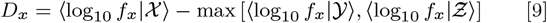

Here 𝒳,𝒴, Ƶ are binary variables that indicate that the input belongs to a specific class. The contrast measures for *y* and *z* outputs are defined analogously.

### C. Evolutionary Training

At each iteration step, we begin with the current best-performing system and generate *M* = 49 of its mutants by evolving the association matrix *K_ij_* according to Eq. (3), with *σ*_mut_= 0.5. For each of these *M* mutants, as well as for the original system, we generate three variants by scaling the concentrations of the hidden layer components by factors 1, 1+Δ, and 1*/*(1+Δ), with Δ = 0.015. To ensure physical plausibility, the evolving association constants and concentrations are constrained by caps: *c*_min_≤ *c* ≤ *c*_max_(*c*_min_= 0.2, *c*_max_= 5.0), and *K_ij_* ≤ *K*_max_= 10_4_*/c*_0_.

Our procedure results in a total of 3(1 + *M* ) = 150 variants that are then evaluated against batches of 48 input cocktails, consisting of 16 cocktails for each of the *n*_out_= 3 output classes. The input concentrations are perturbed by multiplicative log-normal noise: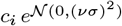. The variant that minimizes the chosen loss function is selected as the new best-performing system, and the procedure is repeated.

### D. Gradient Descent

We start by rewriting Eq.(2) in terms of free concentrations (activities) of the components, *x_i_* = *c_i_f_i_*:

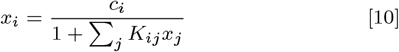

To evaluate gradients of Λ with respect to the model parameters, we expand this equation in terms of small increaments of concentrations, activities and association constants:

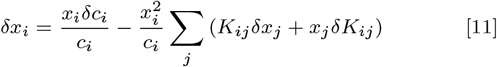

This set of equations can be rewritten as

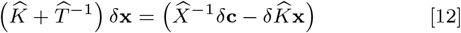

Here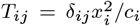and *X_ij_* = *δ_ij_x_i_* This result allows one to calculate the gradients of the loss functions, Eqs. (7), (8), in the space of model parameters,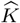 and **c**:

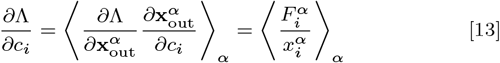

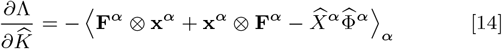

Here:

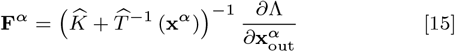

and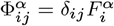. The set of vectors **F^α^** given by given by Eq. 15 can also be evaluated recursively, to avoid inverting matrix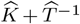at each step, in combination with the recursive solution of Eqs. (1):

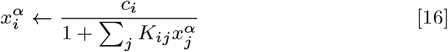

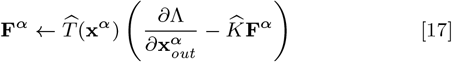

We implemented the gradient descent evolution in terms of concentration of the hidden layer *c*_hid_, and affinity parameters *τ_ij_*≡ ln (*c*_0_*K_ij_* ) = exp(*τ_ij_* ), that are directly related to the corresponding binding free energies: *τ_ij_* = − Δ*G_ij_/RT* . The two loss functions were additionally amended by the regularization term:

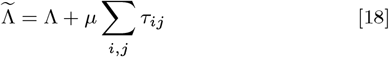

The resulting evolution for the GD evolution of the hidden concentration and the affinity parameters could thus be obtained as

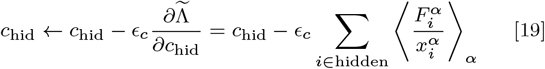

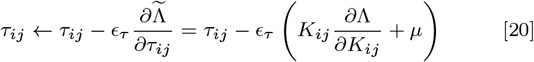

Here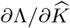and *F* are determined by Eqs.(14),(16), (17), which in turn depend on computing the loss function derivatives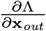:

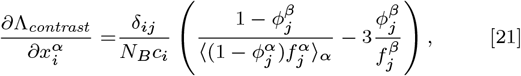

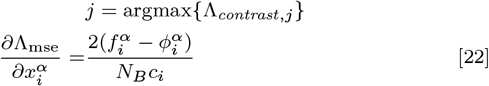

In practice, instead of employing this iterative scheme, we use the adaptive learning rate algorithm from Ref. (28).

### E. Transient behavior

Figures 6-9 illustrate how the loss factions and the performance metric evolve during EL and GD training. It is important to note that each step of EL is highly parallel, as 150 different variants are exposed to 48 different conditions.

**Fig. 6.**
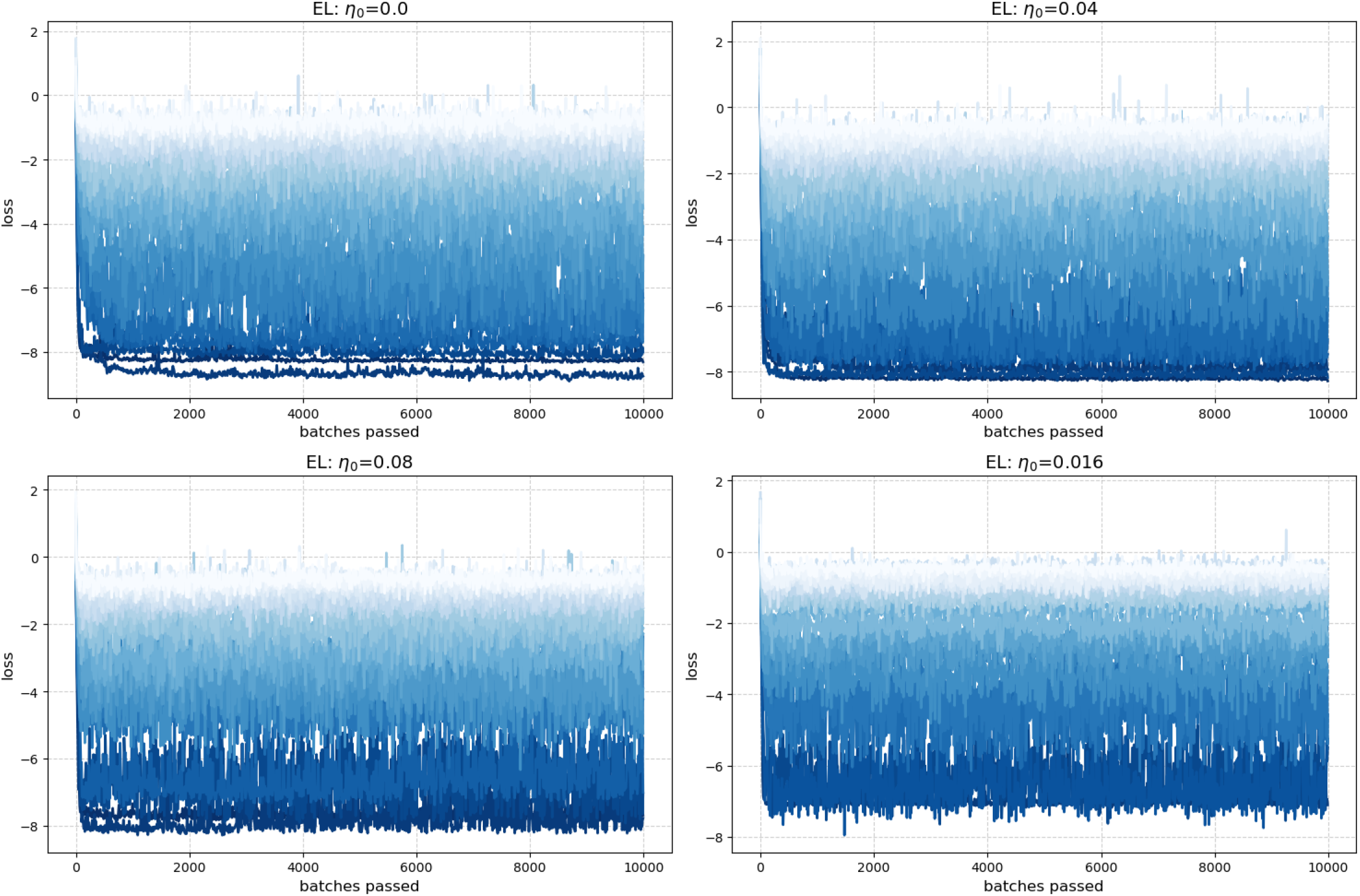
EL: Loss evolution during training for different S/N ration and drift *η*_0_.

**Fig. 7.**
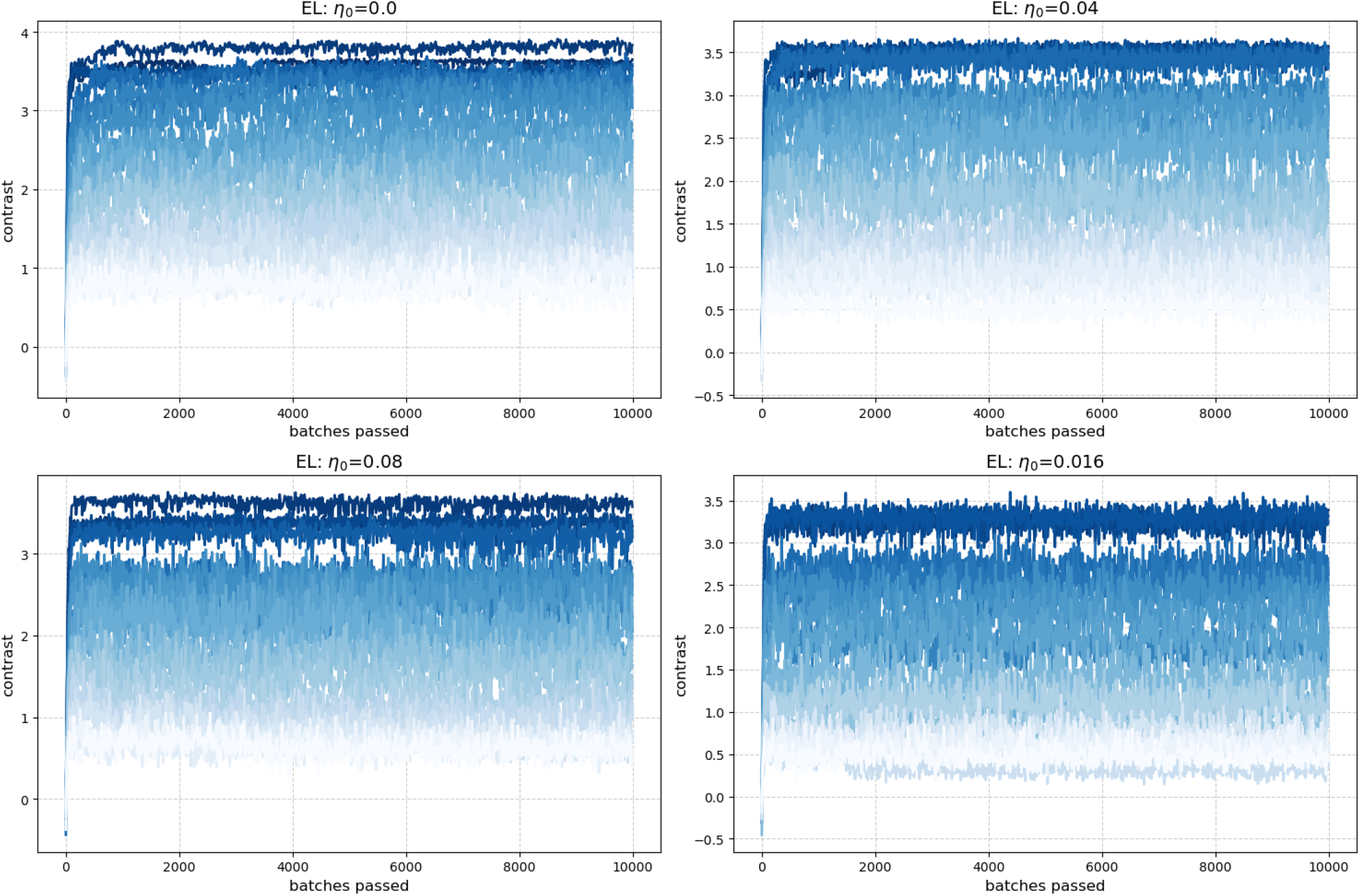
EL: Contrast evolution during training for different S/N ration and drift *η*_0_.

**Fig. 8.**
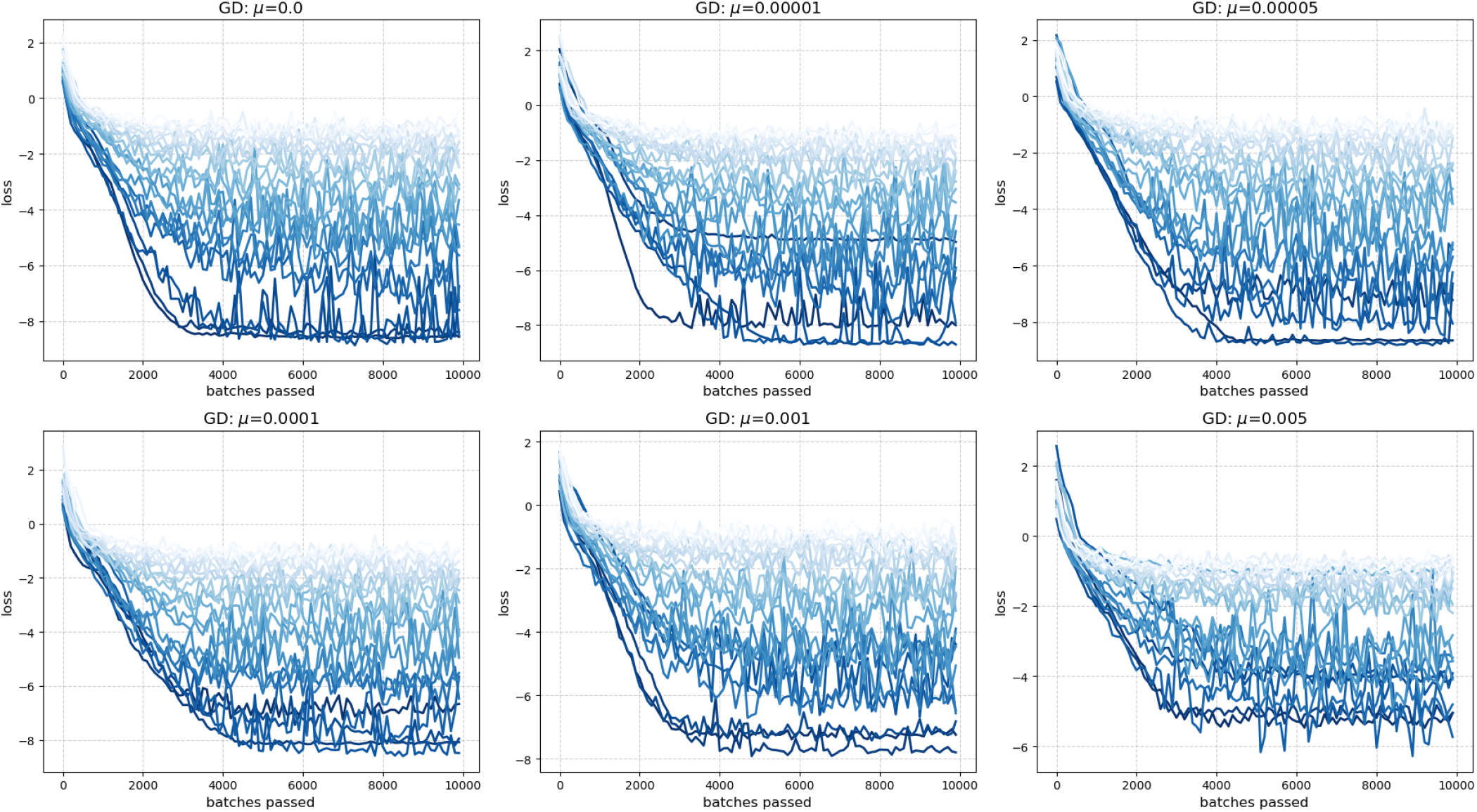
GD: Loss evolution during training for different S/N ration and drift *µ*.

**Fig. 9.**
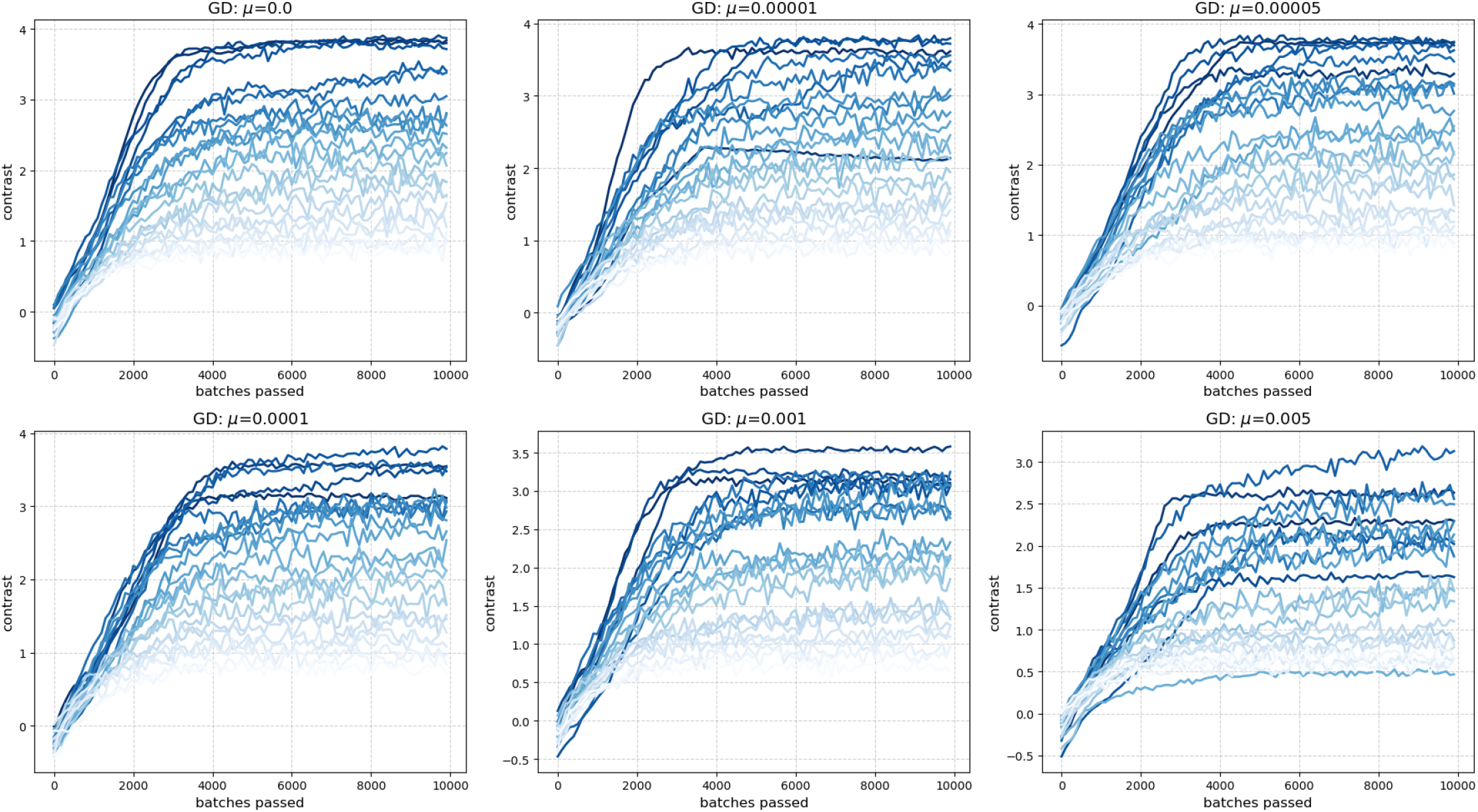
GD: Contrast function evolution during training for different S/N ration and drift *µ*.

## References

1. A Krizhevsky, I Sutskever, GE Hinton, Imagenet classification with deep convolutional neural networks. Commun. ACM 60, 84–90 (2017).

2. I Goodfellow, Y Bengio, A Courville, Deep Learning. (MIT Press), (2016).

3. X Lin, et al., All-optical machine learning using diffractive deep neural networks. Science 361, 1004–1008 (2018).

4. BJ Shastri, et al., Photonics for artificial intelligence and neuromorphic computing. Nat. Photonics 15, 102–114 (2021).

5. C Li, et al., Efficient and self-adaptive in-situ learning in multilayer memristor neural networks. Nat. communications 9, 2385 (2018).

6. A Murugan, Z Zeravcic, MP Brenner, S Leibler, Multifarious assembly mixtures: Systems allowing retrieval of diverse stored structures. Proc. Natl. Acad. Sci. 112, 54–59 (2015).

7. LG Wright, et al., Deep physical neural networks trained with backpropagation. Nature 601, 549–555 (2022).

8. A Momeni, B Rahmani, M Malléjac, P del Hougne, R Fleury, Backpropagation-free training of deep physical neural networks. Science 382, 1297–1303 (2023).

9. M Stern, A Murugan, Learning without neurons in physical systems. Annu. Rev. Condens. Matter Phys. 14, 417–441 (2023).

10. M Stern, D Hexner, JW Rocks, AJ Liu, Supervised learning in physical networks: From machine learning to learning machines. Phys. Rev. X 11, 021045 (2021).

11. M Sitti, Physical intelligence as a new paradigm. Extrem. Mech. Lett. 46, 101340 (2021).

12. D Bray, Protein molecules as computational elements in living cells. Nature 376, 307–312 (1995).

13. US Bhalla, R Iyengar, Emergent properties of networks of biological signaling pathways. Science 283, 381–387 (1999).

14. BA Kramer, J Sarabia Del Castillo, L Pelkmans, Multimodal perception links cellular state to decision-making in single cells. Science 377, 642–648 (2022).

15. W Poole, et al., Chemical Boltzmann Machines. (Springer International Publishing), p. 210–231 (2017).

16. C Floyd, AR Dinner, A Murugan, S Vaikuntanathan, Limits on the computational expressivity of non-equilibrium biophysical processes (2024).

17. S Maslov, I Ispolatov, Propagation of large concentration changes in reversible protein-binding networks. Proc Natl Acad Sci U S A 104, 13655–60 (2007).

18. S Maslov, K Sneppen, I Ispolatov, Spreading out of perturbations in reversible reaction networks. New J Phys 9, 273 (2007).

19. HE Klumpe, J Garcia-Ojalvo, MB Elowitz, YE Antebi, The computational capabilities of many-to-many protein interaction networks. Cell Syst. 14, 430–446 (2023).

20. J Parres-Gold, M Levine, B Emert, A Stuart, MB Elowitz, Contextual computation by competitive protein dimerization networks. Cell 188, 1984–2002.e17 (2025).

21. S Brannetti, S Gentile, ED Grosso, S Otto, F Ricci, Covalent dynamic DNA networks to translate multiple inputs into programmable outputs. J. Am. Chem. Soc. 147, 5755–5763 (2025).

22. MP Nikitin, Non-complementary strand commutation as a fundamental alternative for information processing by DNA and gene regulation. Nat. Chem. 15, 70–82 (2023).

23. TW Hiscock, Adapting machine-learning algorithms to design gene circuits. BMC Bioinforma. 20, 214 (2019).

24. J Shen, F Liu, Y Tu, C Tang, Finding gene network topologies for given biological function with recurrent neural network. Nat. Commun. 12, 3125 (2021).

25. JJ Hopfield, Neural networks and physical systems with emergent collective computational abilities. Proc. Natl. Acad. Sci. 79, 2554–2558 (1982).

26. JJ Hopfield, Neurons with graded response have collective computational properties like those of two-state neurons. Proc. Natl. Acad. Sci. 81, 3088–3092 (1984).

27. J SantaLucia, A unified view of polymer, dumbbell, and oligonucleotide DNA nearest-neighbor thermodynamics. Proc. Natl. Acad. Sci. 95, 1460–1465 (1998).

28. DP Kingma, Adam: A method for stochastic optimization. Preprint: arXiv, 1412.6980 (2014).

